# Ultra-long sequencing for contiguous haplotype resolution of the human immunoglobulin heavy chain locus

**DOI:** 10.1101/2024.12.14.628445

**Authors:** Mari B. Gornitzka, Egil Røsjø, Uddalok Jana, Easton E. Ford, Alan Tourancheau, William D. Lees, Zachary Vanwinkle, Melissa L. Smith, Corey T. Watson, Andreas Lossius

## Abstract

Genetic diversity within the human immunoglobulin heavy chain (IGH) locus influences the expressed antibody repertoire and susceptibility to infectious and autoimmune diseases. However, repetitive sequences and complex structural variation pose significant challenges for large-scale characterization. Here, we introduce a method using Oxford Nanopore ultra-long sequencing and adaptive sampling, coupled with a bioinformatic pipeline, to generate haplotype-resolved single-contig IGH assemblies. We compared our method to a well-established IGH characterization framework using Pacific Biosciences HiFi sequencing in four donors and observed almost complete sequence congruence between our haplotype-resolved assemblies and the HiFi reads. Applying our approach to the HG002 reference material revealed no base differences to the Telomere-to-Telomere genome benchmark over the IGH locus. Importantly, among the four donors, our approach uncovered 30 novel alleles and previously uncharacterized large structural variants, including a 120 kb segmental duplication spanning IGHE to IGHA1 and an expanded seven-copy IGHV3-23 gene haplotype.

## Introduction

Immunoglobulins (Igs) are highly diverse effector molecules integral to adaptive immunity. They play crucial roles in defending against infectious agents but can also contribute to the development of autoimmune diseases. Initially expressed as surface B-cell receptors (BCR), Igs recognize and bind antigens, thereby activating the B cells. This activation drives the differentiation into memory B cells and effector B cells, including plasmablasts and plasma cells, which secrete soluble Igs, or antibodies^1^.

The basic structure of an Ig molecule consists of two identical heavy chains and two identical light chains, forming two heavy-light chain pairs. The Ig heavy chains are encoded within the immunoglobulin heavy chain (IGH) locus, an approximately 1.5 Mb region located at the telomeric end of the long arm of chromosome 14. The locus contains more than 100 homologous gene segments divided into four classes: constant (C), variable (V), diversity (D), and joining (J) genes^2^. The C genes encode the constant region of the antibody, determining the effector functions, while the V, D, and J genes encode the variable domain responsible for antigen binding. The IGH locus exhibits high levels of structural variation (SV) and allelic diversity, contributing to significant variability in the germline repertoire within a population and between the haplotypes in the same individual^3^. During B-cell development, allelic exclusion ensures that only one IGH haplotype is expressed in each B cell. This occurs after successful V(D)J recombination on one chromosome, preventing further recombination on the other chromosome^4^. A highly diverse expressed repertoire is generated through various combinations of V, D, and J gene segments, and additions and deletions at their junctions^5^.

Increasing evidence suggests that variation in the germline IGH locus significantly affects the expressed Ig repertoire^6^. Studies in monozygotic twins indicate that key features in the expressed antibody repertoire are heritable^7,8^, and recent research shows that IGH germline polymorphisms in both coding and non-coding regions influence the V, D, and J gene usage frequencies on a population scale^3^. Furthermore, conserved germline-encoded residues that influence antibody specificity and V, D, and J gene usage have been demonstrated in the immune response against pathogens such as influenza A virus^9^, HIV^10–12^, Staphylococcus aureus^13^, and plasmodium falciparum^14^. Recent studies have also uncovered considerable unexplored germline variation within the IGHC genes^15–17^. Polymorphisms in the IGHC region can influence Fc receptor binding and, thereby, antibody effector functions^18,19^. In membrane-bound IgG, a newly discovered variant in the IGHC region encoding the intracellular IgG1 tail was shown to modulate B cell activation and differentiation, and was also a risk variant for systemic lupus erythematosus^20^. In multiple sclerosis (MS), B cells expressing specific IGHG1 gene variants are selected for in the central nervous system^21^, and correlates with increased intrathecal antibody synthesis^22^.

Our present knowledge of genetic variation within the IGH locus remains incomplete. The high degree of polymorphism, repetitive sequences, CNVs, and SVs have posed significant challenges in characterizing the IGH locus on a large scale^23^. As a result, IG genes have largely been excluded from genome-wide association studies in autoimmune and infectious diseases^24^. Encouragingly, recent technological advancements have enabled the generation of longer sequencing reads with increased accuracy. A framework using capture probes and highly accurate Pacific Biosciences (PacBio) HiFi long-read sequencing allows for high-throughput characterization of the locus, and efforts are underway to gather germline diversity across human populations^25^. However, although this strategy yields highly accurate reads, it may not not necessarily contiguously resolve the entirety of the locus and may miss larger SVs. Oxford Nanopore Technology (ONT) whole-genome sequencing offers increased contiguity of assemblies and improved phasing over longer stretches due to read lengths^26^, but have been hampered by lower raw read accuracy, necessitating deeper sequencing^27^.

Here, we present a protocol that overcomes these challenges without additional library preparation steps before sequencing. We demonstrate that ONT ultra-long sequencing combined with adaptive sampling enables generation of haplotype-resolved *de novo* assemblies of the >1.5 Mb IGH locus. Adaptive sampling, a reference-guided feature unique to ONT sequencing, enables the selective enrichment of regions of interest without prior library targeted enrichment steps^28^. We applied this method to four donors and developed a bioinformatics pipeline to resolve the locus. The results were validated using HiFi reads obtained from the established PacBio single-molecule, real-time (SMRT) sequencing and DNA capture probe method^25^. Our analysis shows great concordance between our assemblies and high-confidence PacBio HiFi reads, with an average of 0.29 nucleotide difference per 1 Mb. Moreover, our method facilitated phasing across larger intervals than previously achieved, consistently resolving all parts of the IGH locus into single-contig haplotypes. This allowed us to resolve novel structural variants, including a large 120 kb segmental duplication spanning the genes IGHE to IGHA1, as well as an expanded 7-copy IGHV3-23 gene haplotype.

## Results

### Generating single-contig haplotype-resolved IGH assemblies

We have established a protocol for comprehensive characterization of the IGH locus using ONT ultra-long sequencing with adaptive sampling. This method was validated on samples from four donors– two healthy individuals (HD1 and HD2) and two MS patients (MS1 and MS2)–as well as the Epstein-Barr virus transformed lymphoblastoid cell line (EBV-LCL) HG002. The donors represent diverse ethnic backgrounds: the healthy donors are of African American and Asian ancestry (HD1 and HD2, respectively) and the MS patients are of European and South Asian ancestry (MS1 and MS2, respectively). We established a sequencing protocol with ONT adaptive sampling, where reads from our region of interest (ROI) were enriched without the need for target amplification during library preparation (Figure 1A). This approach yielded a mean read depth of approximately 30x over the IGH locus (Figure 1B), representing a 5-7 fold increase compared to the mean depth across the genome (Table 1). Furthermore, we obtained high read N50 values ranging from 65-95 kb and read median Phred score between 20.3 and 21.6 (Table 1 and Figure S1).

**Figure 1.**
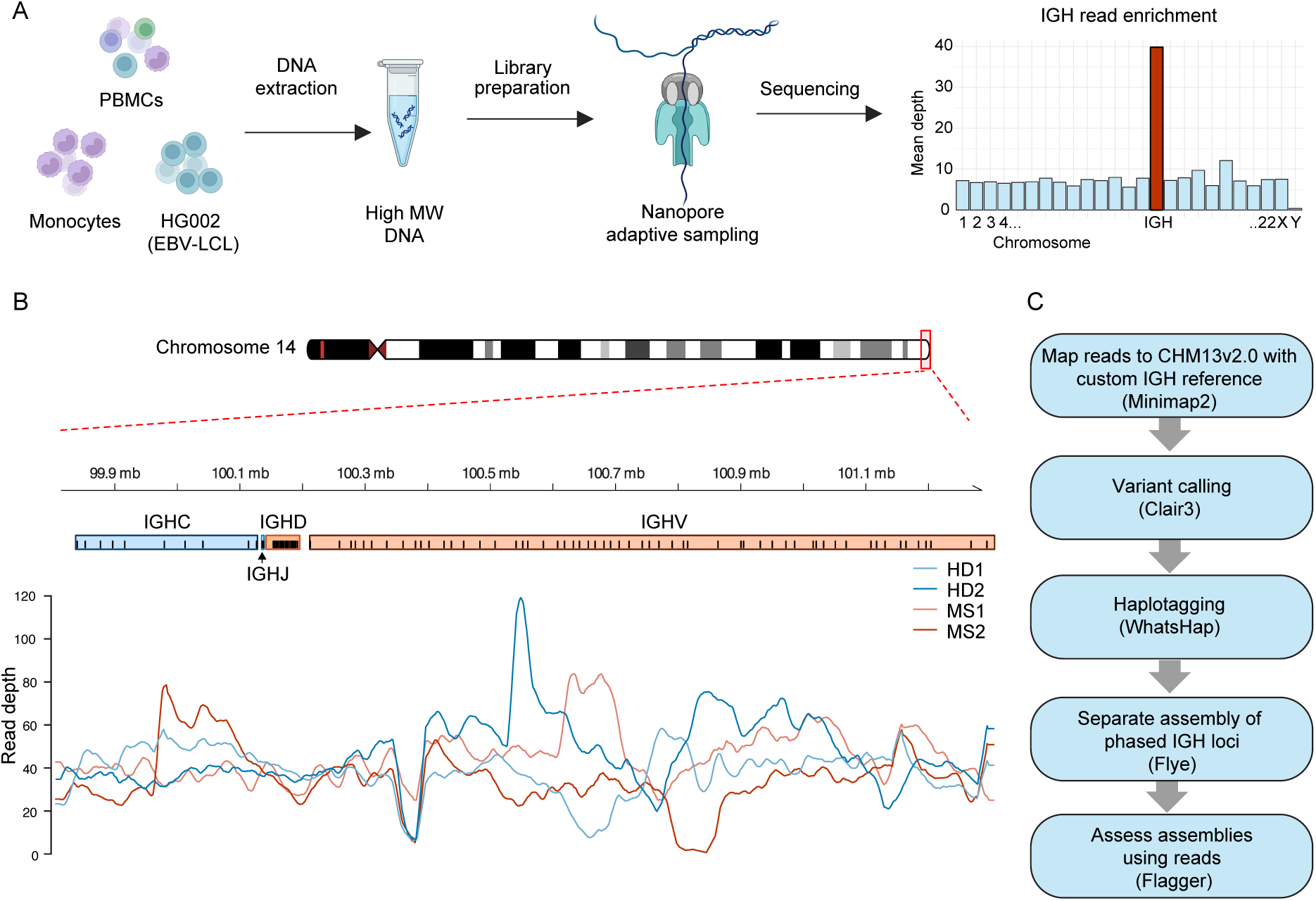
Overview of the experimental setup and bioinformatic pipeline. (A) Schematics of DNA sample preparation and sequencing. High-molecular-weight gDNA was extracted from peripheral blood mononuclear cells (PBMCs), isolated monocytes, or the HG002 Epstein-Barr virus lymphoblastoid cell line (EBV-LCLs). The DNA was used as input for the preparation of ultra long-read Oxford Nanopore sequencing libraries. Sequencing was performed using an adaptive sampling strategy to enrich for reads mapping to the IGH locus. Created in BioRender. Lossius, A. (2024) (B) Read depth from the four donors on a custom IGH reference. (C) Outline of the bioinformatic pipeline used to generate one-contig haplotype-resolved IGH assemblies.

**Table 1.** Descriptive statistics on ONT data input to make IGH assemblies.

| **Individual** | **Longest**<br>**read (bp)** | **N50 (bp)** | **Passed**<br>**reads** | **Mean**<br>**depth** | **Mean IGH**<br>**depth** | **Fold**<br>**enrichment** | **Median**<br>**Phred score** |
| --- | --- | --- | --- | --- | --- | --- | --- |
| **HD1** | 1718964 | 95111 | 335851 | 5.3 | 33.6 | 6.3 | 21.6 |
| **HD2** | 585904 | 83407 | 488896 | 7.0 | 39.8 | 5.7 | 21.0 |
| **MS1** | 944951 | 65367 | 707928 | 6.7 | 40.0 | 6.0 | 20.5 |
| **MS2** | 638907 | 71888 | 403189 | 3.9 | 28 | 7.2 | 20.3 |
| **HG002** | 630829 | 79354 | 566363 | 6.2 | 24.9 | 4.0 | 20.3 |

Using these data, we developed a bioinformatic pipeline employing existing tools (Figure 1C), which enabled the generation of high-quality, contiguous haplotype-resolved assemblies. The pipeline utilizes a custom IGH reference that incorporates several common SVs to enable reference-based phasing of reads. Next, the phased reads are separately assembled *de novo*. We assessed the accuracy of the initial haplotype-resolved assembly drafts, which allowed us to correct phase shifts or revise potential misassemblies (see below). The final corrected IGH loci in all four donors and in HG002 were assembled in single contigs without any apparent phasing ambiguities.

### Assessing the accuracy of IGH assemblies

To ensure the quality of our IGH assemblies, we employed several strategies for rigorous quality control. First, we used the Flagger pipeline to assess assembly reliability^29^. In this approach, reads are mapped back to the corresponding IGH draft assemblies, and coverage distribution is analyzed to identify potential misassemblies. Three of the draft diploid assemblies showed indications of collapsed or erroneous sequences (Figure 2A and Table S1). We verified truly misassembled regions by checking for distinct patterns of SNVs in the read alignments, or by identifying clear boundaries with soft-clipped reads surrounding the region. Segmental duplications pose a challenge for *de novo* assembly due to high sequence similarity between duplicated segments. These SVs are particularly common at this locus, making it crucial to carefully evaluate the assembly and correct any inaccuracies. For instance, one haplotype of the HD1 initially had over 50 kb of its 1.4 Mb IGH assembly flagged as a collapsed region. This was confirmed as a large duplication, which was subsequently corrected. In the final IGH assemblies, only a few bases remain flagged as misassembled, none of which could be confirmed upon manual inspection.

**Figure 2.**
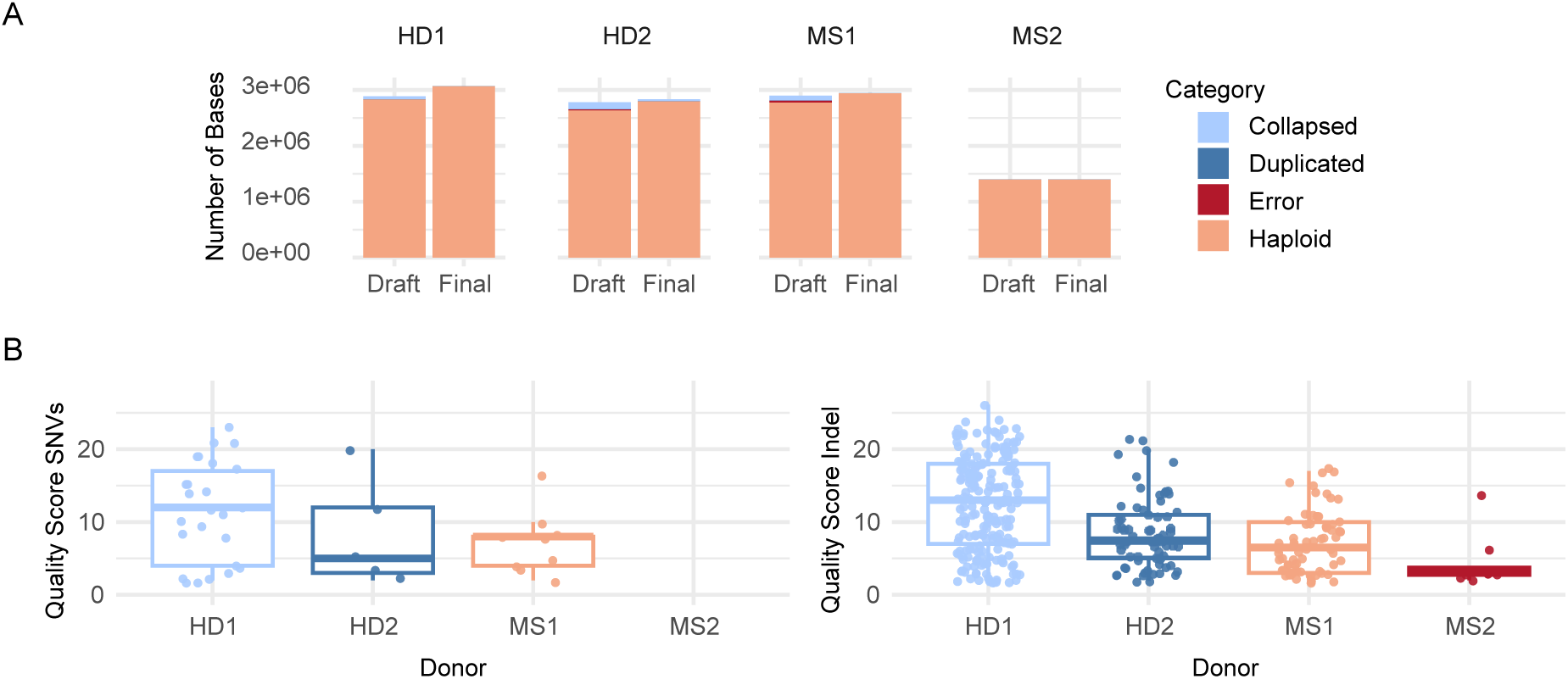
Assessing assembly accuracy. (A) Evaluating assembly accuracy with the Flagger pipeline, where ONT reads are mapped back to IGH assemblies. The column chart illustrates the classification of sequences before and after correction of the diploid assembly. Across assemblies, the sequences are categorized into one of four classes; haploid (indicating a reliable assembly), collapsed, falsely duplicated, or erroneous. (B) Single nucleotide variants (SNVs) and insertions/deletions (indels) identified by PEPPER between the donors’ HiFi reads and their haplotype-resolved IGH assembly from ONT data. Boxplot shows variants were called where read depth was greater than 5x and with quality score greater than 1.

Additionally, we sequenced samples from our four donors using PacBio HiFi technology and used these high-accuracy reads to assess the sequence precision of our ONT assemblies. The HiFi reads were mapped to the final IGH assemblies generated with ONT data, and variants were called using PEPPER with a minimum read depth of 5x (Figure 2B). The HiFi reads showed almost perfect concordance with the IGH assemblies from ONT data. Among the high-confidence variants (Phred score >20), indels were the most prevalent, with only 24 detected across all assemblies. Moreover, only three high-confidence SNVs were detected across the assemblies, which collectively spanned over 10 Mb. Most of these discrepancies were observed in HD1, a unique case that will be discussed further below.

### Benchmarking with HG002

To further evaluate the performance of our long-read ONT method, we sequenced and assembled the IGH locus for the Genome in a bottle (GIAB) HG002 reference material. HG002, also known as NA24385, is a thoroughly sequenced reference genome with extensive publicly available high quality datasets with trio validation. Among these is the recently published telomere-to-telomere (T2T-GIAB Q100) complete diploid HG002 genome^30^. Using our method, we sequenced and assembled the HG002 IGH locus, resulting in one contig per haplotype. These assemblies were very consistent with the Q100 assemblies within the >1.78 Mb encompassing the IGH. In this region, no SNVs were found between the two and only 102 small indels were detected (error rate of 0.000057). When we mapped the Q100 chromosome 14 and our ONT IGH assemblies to the custom IGH reference, we observed as expected that large portions of the HG002 IGH locus and corresponding genes were deleted relative to the custom IGH reference (Figure 3A and B). Notably, our assembly of haplotype 2, corresponding to the paternal haplotype, was missing approximately 900 kb of the ∼1.4 Mb IGH sequence in the reference, including all IGHD genes.

**Figure 3.**
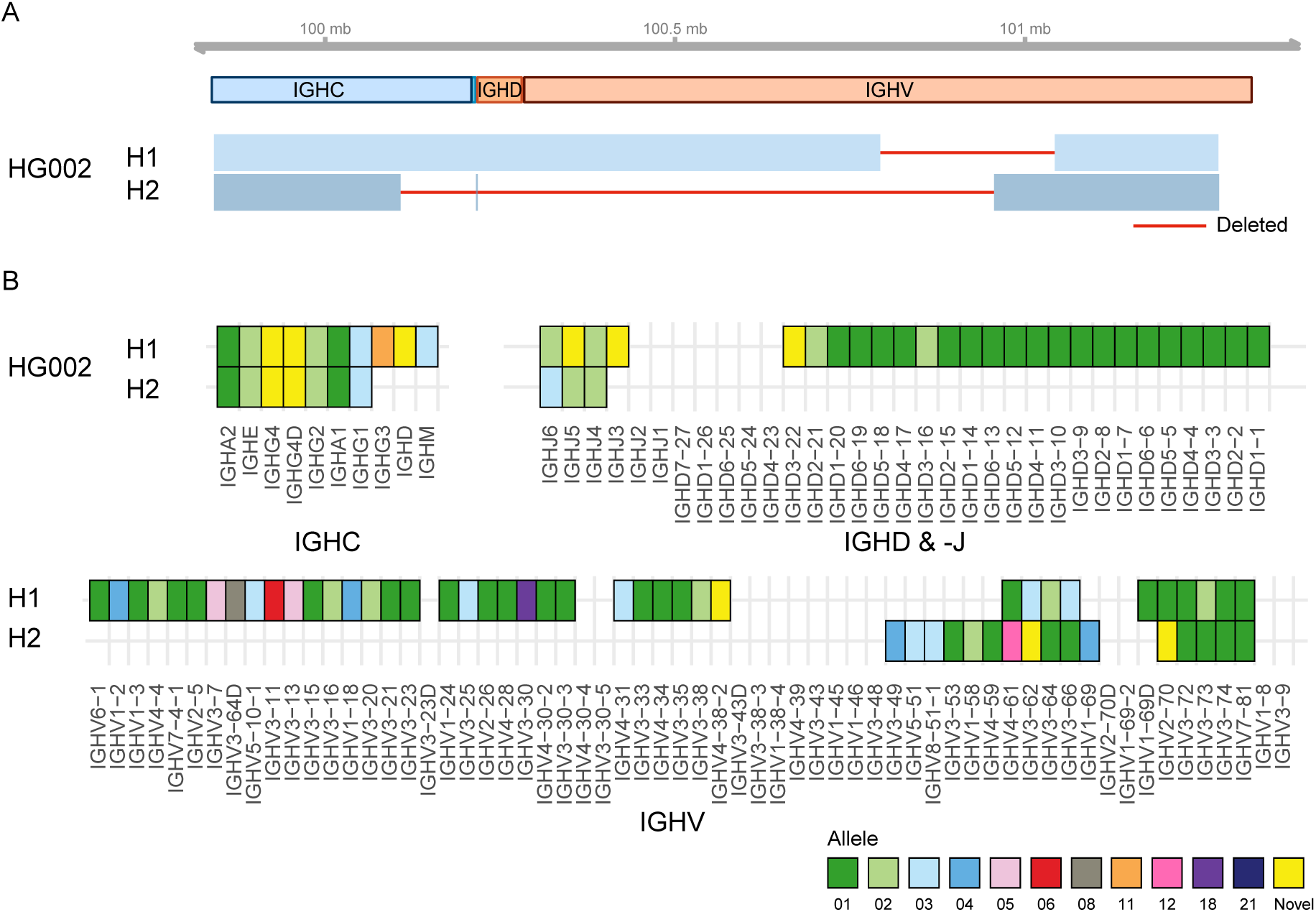
Incompleteness of HG002 IGH assemblies. (A) Schematic representation of the diploid IGH assembly of HG002 aligned to our custom IGH reference. Haplotype 1 (H1) and haplotype 2 (H2) correspond to the maternal and paternal chromosomes, respectively, as derived from the T2T assemblies. The top track displays the IGH gene regions, while the bottom track shows the alignment intervals of HG002 assemblies as blue boxes, with large deletions indicated by solid red lines. (B) Annotation of HG002 IGH assemblies. Empty cell indicates deletion of a gene, and the color of the cell corresponds to the matched allele sequence from the IMGT database. Sequences with no identical match are denoted as “Novel”.

### Annotating assemblies and discovery of novel alleles

A specialized IGH annotation pipeline was applied to the final haplotype-resolved assemblies. Alleles were assigned according to their match identity to known alleles within the ImMunoGeneTics Information System (IMGT) database. Perfect matches to documented IMGT alleles were assigned their corresponding identifiers, while unmatched alleles were classified as “novel” and labeled accordingly. Across the four donor assemblies, we identified a total of 650 V, D, J and C genes, from which we annotated 219 distinct alleles (Figure 4 and Table S2). The majority of these alleles were already characterized and cataloged by IMGT, while 21 of the remaining 51 alleles were found in VDJbase (as of 26.08.2024), and 30 were not found in either database. Notably, among the 69 constant genes annotated in our donors, more than 50% had no corresponding entries in either IMGT or VDJbase.

**Figure 4.**
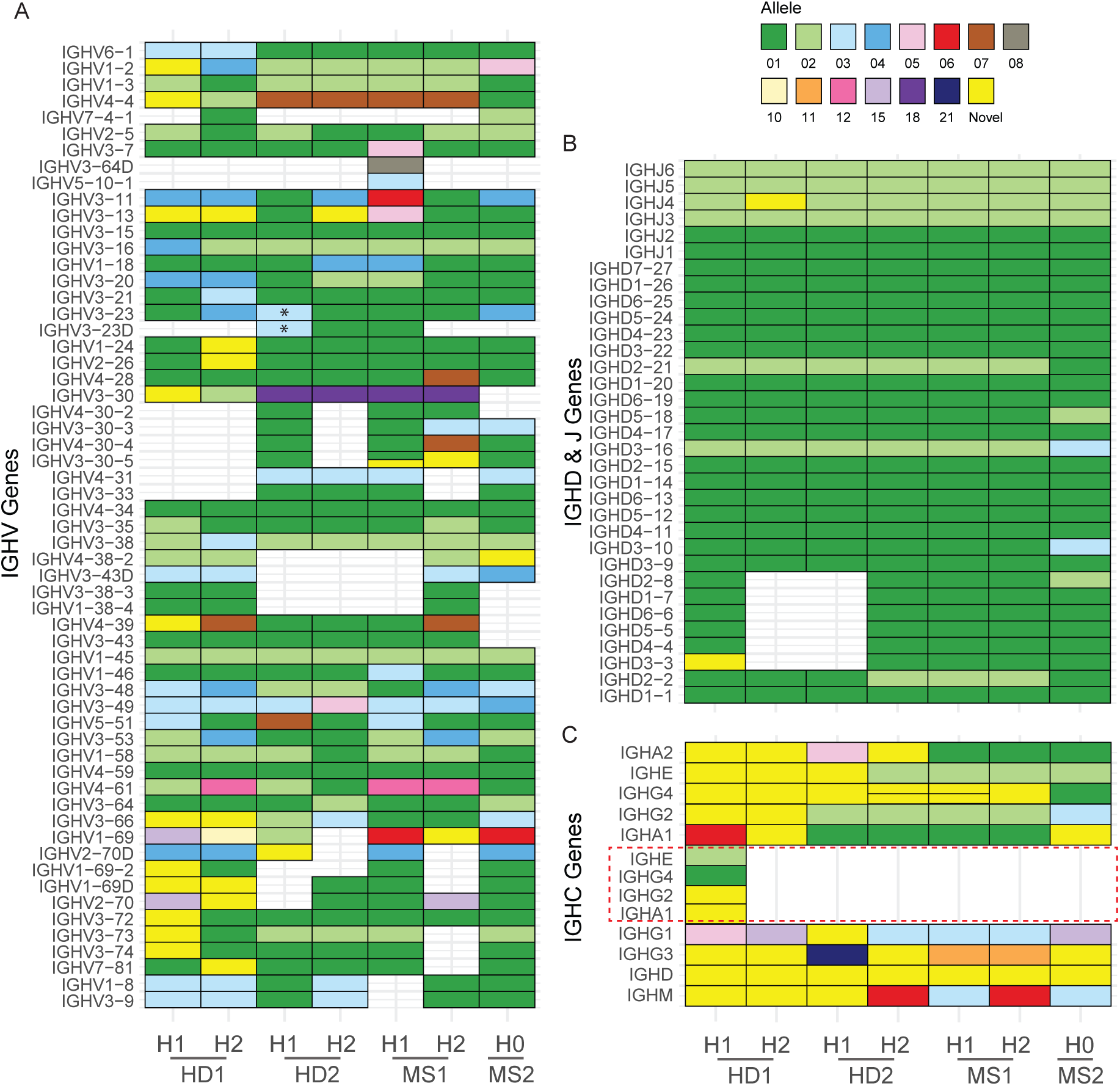
Annotation of IGH assemblies. (A, B and C) Diagrams depicting genes detected in donors’ IGH assemblies. Empty cell indicates deletion of a gene, and a split cell indicates duplication of a gene. The color of the cell corresponds to the matched allele sequence from the IMGT database, sequences with no identical match are denoted as “Novel”. Genes are split by groups, V genes (A), D and J genes (B) and C genes (C). In A, a novel structural variant is denoted by * and represents a large tandem duplication of IGHV3-23 and/or IGHV3-23D, giving a total of seven copies of IGHV3-23. In (C), IGHG4 and its duplicate (sometimes termed IGHG4A or IGHG4D) are joined on one row. The red dotted line indicates a segmental duplication extending from IGHE to IGHA1.

### Novel characterization of a large structural variant in the constant region

A potential large collapse in the IGH constant region was found in one of the draft haplotype assemblies of HD1 (Figure 2A and Table S1), according to Flagger. Inspecting alignment of the donor’s reads to the draft assembly confirmed an increased mean read depth in the flagged region, supporting the presence of a structural anomaly. Further analysis of reads mapping to this region revealed intra-haplotype SNVs with two distinct variant patterns, neither of which completely matched the initial uncorrected assembly. A deeper inspection, involving the alignment of all donor ONT reads to our custom IGH reference, revealed three distinct variant patterns across a ∼100 kb interval in the IGHC region. This finding was corroborated by mapping all HiFi reads to the IGH reference, confirming the presence of three distinct variant patterns spanning the IGHE-IGHA1 interval (Figure 5A).

**Figure 5.**
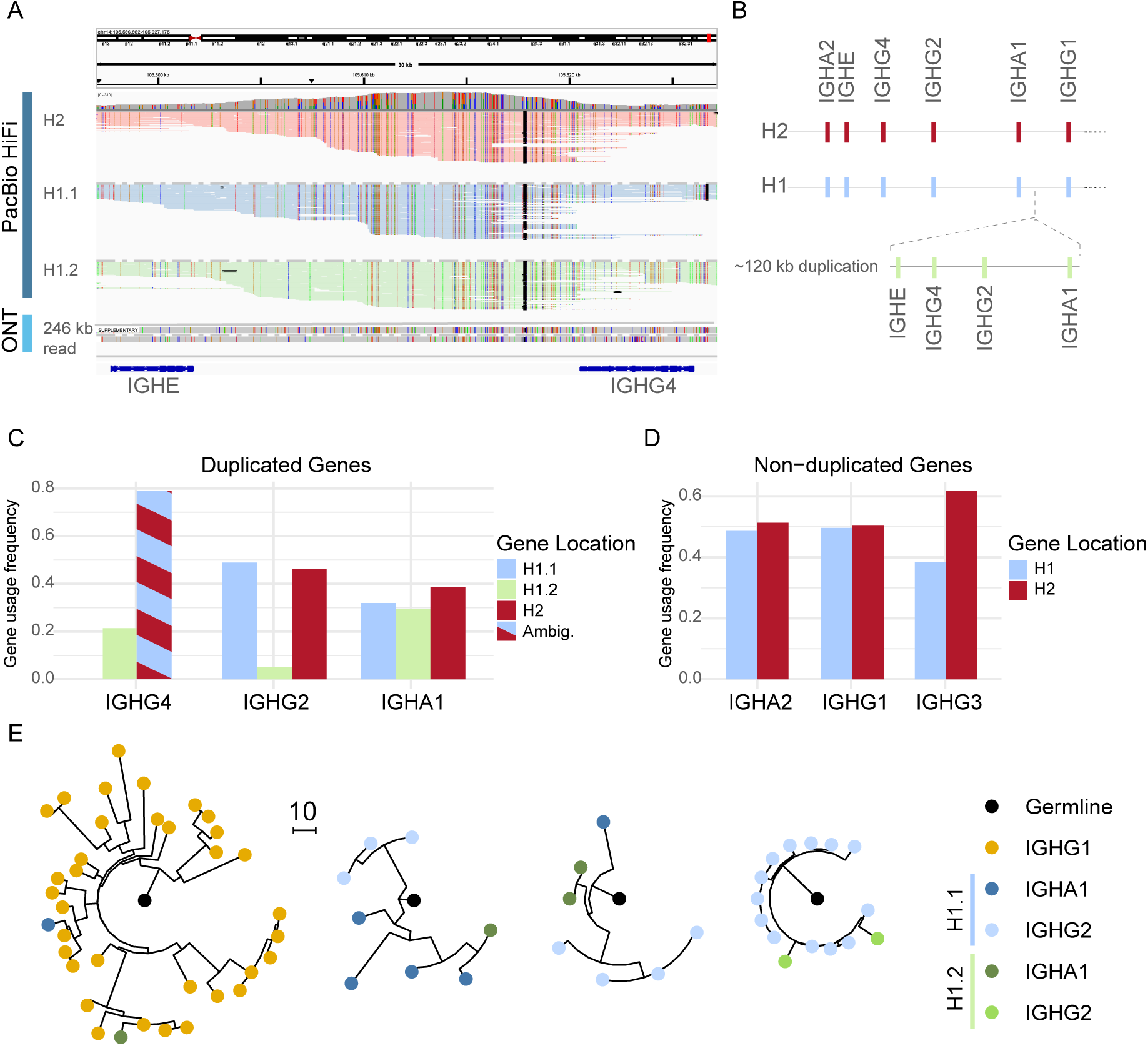
120 kb SV in IGHC. (A) Alignment of PacBio HiFi reads from healthy donor 1 (HD1) onto the hg38 reference genome. The top IGV track shows PacBio HiFi reads grouped by origin: centromeric (H1.1) and telomeric (H1.2) side of the duplication in haplotype 1 (H1), and haplotype 2 (H2). The middle track shows a single ultra-long ONT read (>246 kb) from H1 spanning the duplication, with half of it aligning as a supplementary read. The bottom track illustrates constant gene coordinates. (B) Schematic representation of the large segmental duplication involving four constant genes (IGHE-IGHA1) within HD1 H1. (C and D) Gene usage frequency of each subisotype as determined by FLAIRRseq in HD1. The duplicated genes (C) are stratified by which part of the duplication they originate, except for IGHG4 where the H2 and H1.1 alleles were identical (ambiguous). (D) Frequency of usage of the non-duplicated genes from each haplotype. (E) Representative phylogenetic trees from clonal families containing both copies of duplicated constant genes. Branch lengths indicate mutational distances between nodes, and node colors denote subisotype and gene location.

To further dissect this haplotype, we leveraged our ultra-long ONT reads and identified very long reads (>200 kb) that spanned the reference twice over the IGHE-IGHA1 genes. This strongly suggested that two copies of this array were sequentially arranged on the same chromosome. Collectively, these observations pointed to the existence of a large (∼120 kb) duplication in the IGHC region, which had been collapsed during the *de novo* assembly process. Due to high sequence similarity between the duplicated segments, the variant had to be manually resolved using ultra-long reads spanning the SV, followed by polishing of the assembly with ONT reads. The corrected HD1 haplotype 1 assembly, now including the resolved duplication, showed no conflicts upon realignment of either ONT or HiFi reads (Figure 2A and B, and Table S1).

A proposed model of the SV is presented in Figure 5B. Interestingly, none of the exonic nucleotide sequences of the duplicated genes were identical when compared pairwise. Between the two copies of IGHG2 and IGHG4, there were no coding differences despite three and five exonic SNVs, respectively. IGHA1 contained one nonsynonymous SNV, leading to an amino acid substitution at position 176. The centromeric copy matched IGHA1*06 allele, while the telomeric copy was a novel allele with a D176E mutation. IGHE exhibited the greatest variation among the duplicated genes, with three synonymous and three nonsynonymous SNVs. The telomeric copy corresponded to the IGHE*02 allele, whereas the centromeric copy was a novel allele with mutations A330V, W334G and L464I. This novel allele most closely resembled IGHE*05 from our reference set but differed from the IMGT allele due to the A330V and W334G substitutions.

To determine the extent to which both copies of the duplicated genes were expressed in the BCR repertoire, we performed near-full-length AIRR-seq (FLAIRR-seq)^17^ on PBMCs from the donor. Notably, we identified productive transcripts from both copies of IGHA1 and IGHG2 within the duplicated haplotype 1 (Figure 5C). Since the centromeric copy of IGHG4 on haplotype 1 is identical to that on haplotype 2, calling haplotype for IgG4 transcripts was impossible. The calculated frequency of gene copy usage for each subisotype was evenly distributed between the three copies of IGHA1 (ranging from 29% to 38%). In contrast, IGHG2 transcripts were predominantly derived from the centromeric copy of the gene on haplotype 1 (49%) and the gene on haplotype 2 (46%), while the telomeric IGHG2 on haplotype 1 accounted for only 4.9% of the IGHG2 transcript pool. Finally, we inferred clonal relationships among transcripts from haplotype 1 using SCOPer and identified numerous trees where both copies of a duplicated gene were actively utilized within the same clonal family (example trees are shown in Figure 5E).

### Novel structural variants in the variable region

The donors also harbored several significant novel SVs within the IGHV gene region. In HD2, we identified an expansion of the IGHV3-23/D gene region. Copy number variation has been noted for this region previously^31^, including the recent association with IGHV3-23/D frequency in the expressed Ab repertoire^3^. In HD2, we characterized the first haplotype with seven copies of the IGHV3-23 gene (Figure 6A), the longest expansion so far identified. We extracted the repetitive ∼10.6 kb segments containing IGHV3-23 and compared them to each other (Figure 6B). Interestingly, the five centromeric segments had identical V gene sequence, corresponding to the IGHV3-23*03 allele. The two telomeric copies also shared identical nucleotide sequence corresponding to the IGHV3-23*01 allele. Notably, this donor had two additional copies of IGHV3-23*01 on the other haplotype, resulting in nine diploid copies of this V gene.

**Figure 6.**
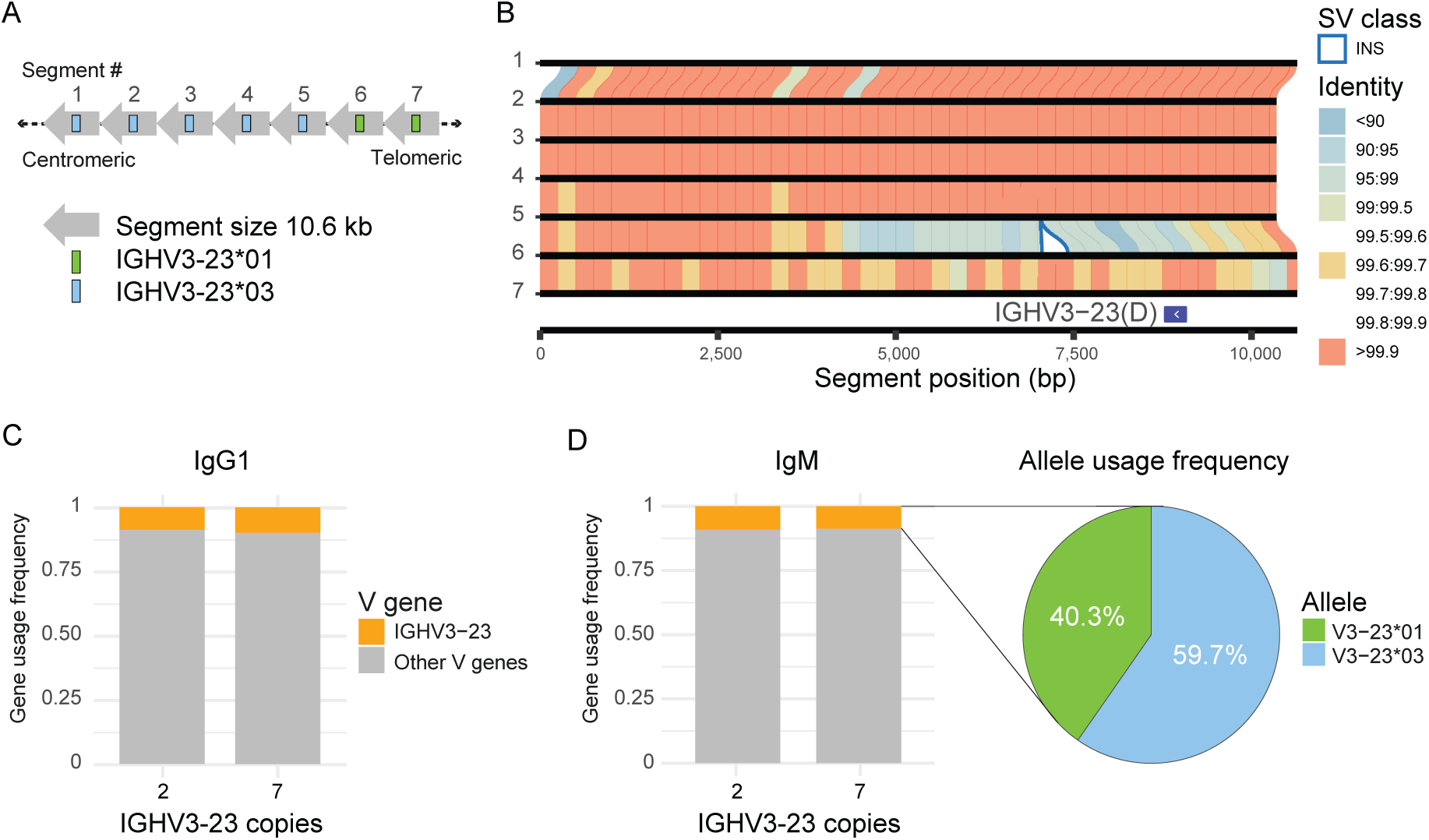
Characterization of a seven-copy IGHV3-23 haplotype. (A) Structural variant giving a seven-copy haplotype of IGHV3-23 in HD2. Illustration shows order of the seven homologous segments. (B) Synteny plot showing sequence identity and structural variation between the seven consecutive IGHV3-23-containing segments. Similarity between segments is represented by lines coloured by % identity of matched bases in 250 bp bins. (C) V gene usage in IgG1 transcripts derived from each haplotype in HD2, a “canonical” two-copy and the novel seven-copy IGHV3-23 haplotype. (D) Frequency of allele usage in IgM transcripts containing IGHV3-23 from the seven-copy haplotype.

To investigate whether the IGHV3-23 gene expansion translated to proportionally higher usage in the expressed BCR repertoire, we performed FLAIRR-seq on PBMCs from the donor. HD2 is heterozygous for IGHG1 and IGHM, which allowed us to differentiate IgG1 and IgM transcripts derived from the two-copy and the seven-copy IGHV3-23 haplotype. Interestingly, our analysis revealed no difference in the proportional usage of IGHV3-23 between the haplotypes, neither in the IgG1 nor IgM repertoires (Figure 6C and D), despite the extra five copies on one haplotype. In the seven-copy haplotype, IgM transcripts utilizing the IGHV3-23 gene were more frequently derived from the *03 allele than the *01 allele (Figure 6D), aligning with the greater number of *03 allele copies.

In addition, HD2 carried single-copy IGHV1-69/IGHV2-70 haplotypes on each of their chromosomes. However, analysis of these haplotypes in the context of the GRCh38 two-copy IGHV1-69/IGHV2-70 haplotype (which harbors, IGHV1-69D, IGHV1-69-2, and IGHV2-70D), indicated that they represent distinct deletion breakpoints. Breakpoint analysis of haplotype 2 was consistent with the deletion of IGHV1−69, IGHV2−70D and IGHV1−69−2 (Figure S2D). However, determination of breakpoint in haplotype 1 was less conclusive as the haplotype is less congruent with the reference (Figure S2C). The haplotype lacks IGHV1-69-2 and IGHV1-69D, but fine mapping of the breakpoint indicates it occurred close to the boundary of - or within IGHV2-70D or IGHV2-70. This means the deletion may not technically be a deletion of one or the other gene, but rather a hybrid of the two. Critically, this is in contrast to the single-copy IGHV1-69/IGHV2-70 haplotype in MS1, which represents the deletion of IGHV1−69D, IGHV1−69−2, and IGHV2−70D, similar to the gene deletion profile previously characterized in GRCh37^3^.

### Hemizygous deletion or identical IGH haplotypes

One of the MS patients (MS2) lacked heterozygous variants necessary for accurate phasing of the IGH locus. Consequently, a non-phased single-haplotype assembly was generated from all reads in the locus. Although a small number of heterozygous variants were detected (Figure S3A), closer examination revealed that these variants had low read support and were located in highly repetitive regions of the locus, consistent with known sequencing errors associated with the ONT platform^32^. The quality distribution of the few heterozygous variants mirrored this observation, in that most variants were low quality (Figure S3B). Furthermore, when corresponding HiFi reads were analyzed, no high-confidence SNV or indel calls were identified when mapped to the haploid assembly (Figure 3B). These findings suggest that this individual is either hemizygous for the IGH locus or possesses two identical haplotypes. Accordingly, further bioinformatic separation of the two haplotypes was not warranted.

## Discussion

In this study, we present a method that utilizes ONT ultra-long sequencing with adaptive sampling to generate phased, accurate and complete IGH assemblies without the need for complementary support by other technologies. By employing DNA extraction techniques to preserve ultra-long reads, we achieve an N50 exceeding 65 kb, with some reads extending beyond several hundred kb. This is complemented by a bioinformatics framework that assembles, phases, quality-controls, and comprehensively annotates the entire IGH locus. The value of the method is demonstrated by its ability to uncover significant structural variation even in a very limited cohort.

Among these findings is a ∼120 kb duplication in the IGHC region, which to our knowledge is the first sequence-level description of this variant. More than 30 years ago, a duplication in the same region was reported in an Italian family using Southern blot analysis^33^. However, it is not possible to determine whether this duplication corresponds to the same haplotype or represents an individually arising variant. To this point, such large homologous duplications would not have been possible to resolve using the short-read technology at the time and challenging to discover serologically when relying on sparse coding variation. Importantly, the variant appears to have functional consequences for the expressed B cell receptor repertoire, as the sequencing of IgA and IgG transcripts from the donor showed productive transcripts containing all three copies of IGHA1 and IGHG2 (Figure 5C). Interestingly, the gene usage frequency varied significantly between the isotype groups. All copies of IGHA1 were used similarly, which meant that 61.5% of the IGHA1-containing transcripts came from the duplicated haplotype. In contrast, less than 5% of the IGHG2 transcripts contained the haplotype 1 telomeric copy.

We have demonstrated that all duplicated constant genes are participating in class-switch recombination. However, we also wanted to know if both copies were utilized within the same clonal family. Indeed, we found multiple instances of clonal families employing both copies of the duplicated constant genes. This observation suggests the possibility of non-canonical class switching occurring between the duplicated copies, allowing, for example, a B cell to switch from IGHG4 to IGHA1. Alternatively, the co-occurrence of the duplicated genes within a family could be explained by clonally expanding B cells class switching to different copies of the duplicated gene separately.

A large tandem duplication expansion encompassing IGHV3-23 was found in the healthy donor HD2. This haplotype had seven copies of the IGHV3-23 gene, wherein five consecutive segments contained the *03 allele and the last two were *01 (Figure 6A). The five ∼10.6 kb duplicated segments containing the *03 allele displayed almost identical sequences even beyond the V gene exon (Figure 6B), which suggests that in some multi-copy IGHV3-23/D haplotypes will require long reads (>50 kb) to correctly resolve copy number variation. However, designation of these duplication blocks to those harboring IGHV3-23 or IGHV3-23D among the 2-copy is non-trivial, suggesting that analysis of additional multi-copy haplotypes from diverse populations will likely be required to effectively characterize gene copy number, allelic polymorphisms, and non-coding SNVs.

We were interested in the effect of this expansion on the expressed repertoire, as HD2 carried a total of nine diploid copies of the IGHV3-23 gene. IGHV3-23 is associated with influenza response^34^ and is one of the most used variable genes in the BCR repertoire^35^. It has been shown that copy number variation of IGHV3-23 has a significant additive effect on its gene usage^3^. Therefore, it was surprising to find that IGHV3-23 usage fell within the range of what has previously been published, and that there was no difference in gene usage between the haplotypes despite a 3.5 fold difference in gene copy number between them.

Interesting deletion haplotypes were also present in our donor cohort. Closer analysis of putative deletion breakpoints among characterized single-copy IGHV1-69/2-70 gene haplotypes indicates that this region is likely the site of recurrent SVs, and that there are at least two deletion-haplotypes circulating in the human population. These deletion haplotypes, in addition to the other SVs identified here, highlight considerations for the characterization and cataloging of both coding and non-coding variation in these regions, in particular as this pertains to the naming and curation of gene allelic variation. For example, with respect to the IGHV1-69/D and IGHV2-70/D genes, alleles from single copy haplotypes have been historically assigned to IGHV1-69 and IGHV2-70. However, our analysis shows that the accurate allele assignment will require more nuanced and high-resolution analysis of gene copy number and haplotype variation to make such designations. Likewise, it will be non-trivial to name and assign alleles to the new IGHV3-23 and IGHC genes identified in this study, and highlights that we should proceed with caution to do this robustly.

Existing databases are far from covering the full extent of allelic polymorphism in Ig genes within the human population^3,36^. In our study, we analyzed samples from two MS patients and two healthy donors, identifying 25 novel IGHV alleles and 25 novel IGHC alleles not present in the IMGT database. These novel alleles were independently validated using mapped HiFi reads, with each allele supported by at least ten HiFi reads. Compared to allele inference using adaptive immune receptor repertoire sequencing (AIRR-seq), a major advantage of our method is its ability to detect variants in the non-coding regions of IGH. This is in the same vein as the established PacBio SMRT-seq and DNA capture probe method, which was recently used to explore the impact of germline variation on the expressed repertoire^3^. The study revealed that SNVs associated with variation in IGHV gene usage were enriched in intergenic regions involved in VDJ recombination, thereby explaining previous observations of biased allele usage inferred from AIRR-seq data^37^.

Variant calling during phasing revealed a strikingly different variant profile between the donors. This was particularly pronounced in one of the MS patients (MS2), where we observed a significant skewing towards homozygous variants, leading to poor haplotagging resolution. Accurate read phasing relies on an even and frequent distribution of high-quality heterozygous variants; consequently, MS2 had unusually small phase sets due to the scarcity of such variants. This strongly suggests that the individual may have two identical haplotypes or could be hemizygous for the IGH locus. While terminal 14q deletions have been reported, they are rare and typically associated with cognitive disability and dysmorphic features^38,39^, neither of which were present in our patient, who was otherwise healthy apart from MS. Moreover, the patient had normal serum levels of IgA, IgM, and IgG (data not shown) and no history of increased susceptibility to infection. Additional investigations, such as *in situ* hybridization or qPCR, could have provided further insights but were considered outside the scope of this study.

We observed significant deletions in both maternal and paternal haplotypes of the HG002 EBV-LCL at the IGH locus. EBV-LCLs are established by infecting a heterogeneous pool of B cells with EBV, resulting in the creation of immortalized cell lines. The mature B cells targeted during this process have undergone recombination of their IG loci. Therefore, depending on the clonality of the transformed B cell pool that makes up the EBV-LCL, substantial portions of the original germline IGH locus are expected to be absent^40^. The clonality of our HG002 cell line is unknown, but as the IGH assemblies resolved without ambiguity and nearly identically to the T2T-GIAB Q100 assemblies we should assume clonality is low if not monoclonal. This aligns with a recent preprint, which found a low number of recombination events in HG002 using data from the Human Pangenome Reference Consortium^41^.

Acquisition of diploid IGH assemblies represent a significant advancement in understanding the structure and regulation of the IGH locus. VDJ recombination and class-switch recombination occur on a single chromosome and involve cis-regulatory elements that can function over a distance of several 100 kb^42^. Polymorphisms within these regulatory elements can lead to changes in epigenetic regulation, thereby impacting the formation of the immune repertoire^43^. The mechanisms may involve interactions with transcription factors and other regulatory proteins, and/or alterations in locus topology^44^. Diploid assemblies are essential to dissect these complex mechanisms, which can be achieved by combining long-read sequencing technology with epigenetic assays. Since a given B cell clone expresses one of the two haplotypes, diploid assemblies are also valuable for analyzing the clonal evolution of B cells at the level of IGHC alleles^17^. Such approaches can link VDJ and IGHC genetic signatures, providing new insights into the interaction between the Ig constant and variable regions.

Our method successfully assembled all IGH haplotypes from four individuals and one cell line into single phase blocks and resolved novel structural variants. To achieve this, we had to optimize several key steps in our pipeline. Firstly, to ensure consistent results across all individuals, we experimented with various assembly and phasing strategies. For instance, creating a consensus haploid assembly as a starting point before converting it into a diploid assembly, as suggested by others^26^, often led to misassemblies if there were large SVs between the two haplotypes.

Consequently, we found that a reference-based phasing strategy produced the best overall phasing and assemblies for all individuals. Secondly, an essential component of our pipeline is the quality control of the assemblies, which involves mapping reads back to the assembly draft in a haplotype-aware manner. Misassemblies were detected using Flagger, which estimates the expected read coverage and flags regions with coverage inconsistencies that are likely to be assembly errors^29^. This enabled us to identify issues such as the large duplication in the constant region in HD1, which required manual curation. The historically low read accuracy of ONT reads has previously posed a challenge for using the technology independently, often necessitating the combination with short-read technologies to enhance accuracy^45^. However, recent advancements of the ONT platform have substantially improved its sequencing accuracy^32^. In our present pipeline, we utilized R10.4 flow cells in conjunction with the latest super-accurate base-calling mode, achieving an estimated mean read accuracy exceeding 99%. With a diploid read depth of approximately 30X over the IGH locus, this high accuracy translated into near 100% accurate assemblies. Accordingly, we observed a near-complete sequence concordance with HiFi reads from an established probe-based framework^25^, and benchmarking our method using the HG002 reference genome revealed no SNVs compared with the Q100 published genome. This level of accuracy, previously unattainable with ONT alone, signifies a milestone in long-read sequencing, enabling highly accurate assemblies without the need for short-read polishing.

Currently, our method has some technical limitations that reduce its scalability for large cohorts. Firstly, the adaptive sampling technology is quite permissible for reads outside the ROI, resulting in reduced sequencing capacity due to pore occupancy by off-target reads. However, with further improvements of this technology, patients could be multiplexed on the same flow cell and the protocol expanded to simultaneously enrich other immune loci like IG light chains, TR or HLA. This would allow for a more comprehensive characterization of germline immune genetics and open up for holistic approaches into investigating germline variation effects on adaptive immune repertoires. Secondly, the bioinformatic tools for phasing and assembly are not yet fully optimized for the complexities of the IGH locus, so manual inspection and correction of intermediary files are still necessary, making the process labor-intensive. With the availability of larger datasets of IGH haplotypes in the future, construction of a panlocus reference graphs and variant calling with this could eventually eliminate the need for *de novo* assembly entirely.

Nevertheless, the method described in this paper provides substantial utility for characterizing the IGH locus in both health and disease contexts—a task that has historically been challenging due to the limitations of short-read sequencing in resolving intricate genetic sequences and structures. This approach opens new avenues for exploring genetic diversity and commonalities across populations and patient cohorts, facilitating the discovery of inherited genetic patterns that influence adaptive immune responses.

## Materials and methods

### Ethics statement

The study and sample collection was approved by the Regional Ethical Committee South East Norway (2009/23) and the UofL Institutional Review Board (IRB 14.0661). All donors provided written informed consent before study inclusion.

### Sample collection and preparation for ONT sequencing

Peripheral blood mononuclear cells (PBMCs) were collected from two MS patients using Vacutainer CPT (Becton Dickinson Biosciences), before monocytes were negatively selected using the Pan Monocyte Isolation Kit, human (Miltenyi Biotec). Cells were frozen and kept in nitrogen until DNA extraction and sequencing as stated below.

Commercial frozen human PBMCs were purchased from StemCell Technologies. HG002 EBV-LCL were purchased from Coriell Institute of Medical Research and expanded for 14 days in Gibco RPMI 1640 (Thermo Fisher Scientific) supplemented with 15% heat-inactivated fetal calf serum and Penicillin/Streptomycin at 100 U/ml and 100 ug/ml, respectively, before freezing aliquots.

### ONT DNA library preparation

Approximately 6 x 10^6^ cells, either monocytes (MS1 and MS2), PBMCs (HD1 and HD2) or EBV-LCLs (HG002), were used as input for DNA extraction and subsequent ultra-long sequencing library preparation. DNA was extracted using Monarch® HMW DNA Extraction from Cell & Blood (New England Biolabs) following the protocol “High Molecular Weight DNA (HMW DNA) Extraction from Cells (NEB #T3050)” with the following alterations: 1. The agitation speed during cell lysis was set to 1400 rpm for 10 minutes, 2. Incubation time with beads during gDNA binding to beads was increased to 8 minutes, 3. Elution buffer was replaced with 560 µl Extraction EB from SQK-ULK114 (ONT), 4. An extra incubation step for 1 hr at room temperature was added after elution with agitation at 56°C, 5. The centrifugation of beads with DNA was increased to 1 minute at 16,000 x g, 6. An additional 200 µl Extraction EB buffer was added after complete elution to get correct input volume for ONT library preparation. Sequencing libraries were generated using Ultra-Long DNA Sequencing Kit from ONT (SQK-ULK114) following the manufacturer’s guidelines.

### ONT sequencing

Sequencing was performed on a PromethION 2 solo (P2 solo) device connected to a high-performance computer running MinKNOW software version 23.07.12. Loading and washing of PromethION R10.4.1 flow cells was performed as per manufacturer’s guidelines. Each patient was sequenced using one flow cell and the sequencing ran for a maximum of 72 hrs with washing and reloading performed around every 24 hr using the flow cell wash kit (EXP-WSH004) to increase output.

To serve as an IGH reference for adaptive sampling during sequencing, the T2T-CHM13v2.0 was altered at the telomere end of chromosome 14. Here, the variable part of IGH was replaced with a custom IGH reference as described elsewhere^25^ whilst the rest of chr14 including the IGH constant region remained. Adaptive sampling was enabled in MinKNOW to enrich for the ROI, specified as the IGH region with an additional 200 kb lead-in from the custom CHM13 reference. During sequencing, the high accuracy basecalling model by Dorado (v 7.1.4) was enabled to accommodate adaptive sampling. The resulting POD5 files were rebasecalled post-run using the super-accurate model, filtering out reads with a Phred score below 10, and subsequently used for further downstream processing.

### Processing ONT data to IGH assemblies

A pipeline to process ONT reads into haplotype-resolved IGH assemblies was established using existing tools and custom scripts. Firstly, catfishq (version 1.4.0) was used to filter out adaptive sampling rejected reads from the rebasecalled FASTQ files. Next, the reads were aligned to our CHM13v2.0 reference with custom IGH described above using Minimap2^46^ (v 2.26) with option ‘-ax map-ont’. Variant calling was performed on the resulting BAM file with Clair3^47^ (v 1.0.0) with options ‘--enable_phasing’, ‘--platform=’ont’’ and ‘--bed_fn’ enabled specifying a BED file with IGH coordinates (chr14:99630000-101325184). Reads were haplotagged using Clair3 <phased_merged_output.vcf> with Whatshap^48^ (v 1.4) haplotag and options ‘--output-haplotag-list -- ignore-read-groups --tag-supplementary --region=chr14:99630000-101325184’. Continuity of haplotype tags between phase sets was evaluated by manual inspection of the haplotagged BAM in the Integrative Genomics Viewer (IGV)^49^. If switch errors were detected, the Clair3 VCF was edited before running ‘whatshap split‘ with default parameters to produce hap1 and hap2 phased FASTQs. Lastly, *de novo* assembly was produced from the haplotype-separated reads using Flye^50^ (v 2.9.2), with arguments ‘--genome-size 1200K --nano-hq <phased.fastq>’.

Manual inspection in IGV and Flagger^51^ (v 0.4.0) was used to evaluate haplotype-resolved assemblies. ONT reads from the ROI were mapped back to the diploid assembly with Minimap2 and Flagger highlighted misassembled regions. Problematic regions of the assemblies had to be manually resolved when collapsed by the assembler. In the example of the collapsed large segmental duplication, we isolated reads that spanned the duplication. Subsequently, one of the longer isolated reads was extracted to a fasta and polished with the rest of the isolated reads using ‘flye --nano-hq <isolated_reads.fastq> --polish-target <long_read.fasta> --iterations 3’. The polished long read was mapped back to the first-draft assembly with Minimap2 with ‘-ax asm20’ option and coordinates for cutting and pasting of the resolved duplication were determined. The entire collapsed sequence was removed and the polished long read was trimmed to fit this gap in the assembly and inserted here. The resulting new draft assembly was finally verified again as outlined above with Flagger.

### PacBio HiFi sequencing of IGH using capture probes

Parallel sequencing of samples from all donors was performed according to the previously published method^25^. In brief, probe-based targeted capture of the IGH region and long-read SMRT-seq was used to sequence IGH reads. Libraries were sequenced on PacBio instruments SEQUEL II (HD1, MS1 and MS2) or REVIO (HD2).

### IgA and IgG FLAIRRseq

RNA was extracted from HD1 and HD2 PBMCs using AllPrep DNA/RNA Mini Kit (Qiagen). IgG and IgA transcripts were resolved by targeted amplification using near-full-length Adaptive Immune Receptor Repertoire sequencing (FLAIRR-seq) as described previously^17^. FLAIRR-seq cDNA libraries were prepared using the SMRTbell Prep Kit 3.0 (Pacific Biosciences) according to the manufacturer’s specifications. The prepared library was sequenced on the Revio long-read system (Pacific Biosciences). FLAIRR-seq data was processed via the Immcantation tool suite using pRESTO and Change-O as previously described^17,52,53^. For the identification of clonal relationships, we used SCOPer hierarchical clustering with a threshold of 0.15^54^. Dowser and Igphyml were used to build and visualize the clonal trees^55,56^.

### Validation of IGH assemblies

Personalized alignment of HiFi long reads were used to infer orthogonal validation and base level concordance was obtained for ensuring the highest quality of haplotype resolved assemblies. The HiFi reads were aligned with Minimap2 to the corresponding ONT IGH assemblies with option ‘-ax map-hifi’. The variant caller function of PEPPER-Margin-DeepVariant (v 0.8) was used to assess base accuracy as a measure of agreement between the high-accuracy reads and the polished assemblies. PEPPER-deepvariant image was ran with singularity and the options ‘call_variant -- pepper_min_coverage_threshold 5 --hifi’ The alignment of HiFi reads were also used to infer orthogonal validation and base level concordance was obtained with other metrics. Custom Python scripts were utilized to assess the proportion of assembly bases with at least 80% concordance from the reads, ensuring a minimum read depth of 5X and 10X for estimating coverage and accuracy.

### Annotating IGH assemblies

IGH assemblies were annotated in a reference guided approach using custom python and bash scripts. Finished ONT assemblies and reads were aligned to a human reference genome^25^ using Minimap2 with presets ‘-ax asm20’ and ‘-ax map-ont’, respectively. Sequences for IGH exons were extracted from assembly bam using gene coordinates mapped to the reference from GRCh38 and searched against available human IGH alleles in IMGT database (IGHV, -D and -J sets were downloaded February 13th 2024, IGHC alleles collected November 24th, 2022). HiFi reads mapping over the same intervals were used to validate support for assigned and novel alleles, using the same python scripts and metrics that were used to assess read coverage in the assembly as a whole.

## Acknowledgements

We express our sincere gratitude to the patients and donors whose contributions were vital to this study. We wish to thank Steffan Daniel Bos-Haugen for useful discussions in the initiation of this project, Milda Kaniušaitė for her valuable guidance on the wet lab and sequencing protocol, and professor Tone Tønjum for providing access to sequencing equipment during the initial phase of the study. This research was supported by grants from the Norwegian Research Council (project no. 314376) and the South-Eastern Norway Regional Health Authority (project no. 2021085).

## Author contributions

A.L., E.R. and C.T.W. contributed to the conception of the work. M.B.G., E.R., and E.E.F. performed the sequencing. M.B.G., A.L., A.T., W.D.L., Z.V., U.J. and C.T.W. developed the computational pipeline to analyze the ONT sequencing data. M.B.G., A.L., U.J., A.T., C.T.W. and E.E.F. analyzed sequencing data from ONT, PacBio and/or FLAIRR-seq. A.L., E.R., M.L.S., and C.T.W. acquired funding. M.B.G. and A.L. drafted the manuscript. All authors contributed to revising the manuscript.

## Declaration of generative AI in the writing process

The initial draft was written without the use of any generative AI tools. In some instances, the authors employed ChatGPT to enhance the wording of a sentence. In these cases, the authors reviewed and edited the content as needed and take full responsibility for the content in the publication.

## Data availability

Raw reads from ONT sequencing of EBV LCL HG002 will be available on SRA at the date of publication: https://www.ncbi.nlm.nih.gov/sra/PRJNA1196145

Raw sequencing reads from ONT and PacBio platform, sourced from MS patients and donors, are currently being uploaded to the European Genome-Phenome Archive (EGA).

