## Supplementary Figures and Tables for "Ultra-long sequencing for contiguous haplotype resolution of the human immunoglobulin heavy chain locus"

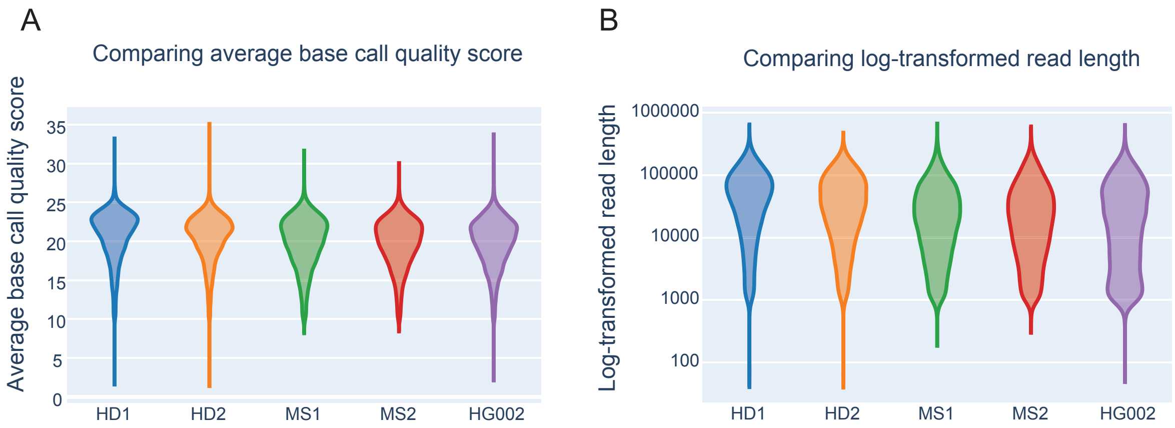


**Figure S1. Descriptive statistics on all rebasecalled sequencing reads, excluding those rejected by adaptive sampling**. (A) Violin plot of base call quality scores from all donors and HG002. (B) Log-transformed violin plot of read lengths from all donors and HG002.

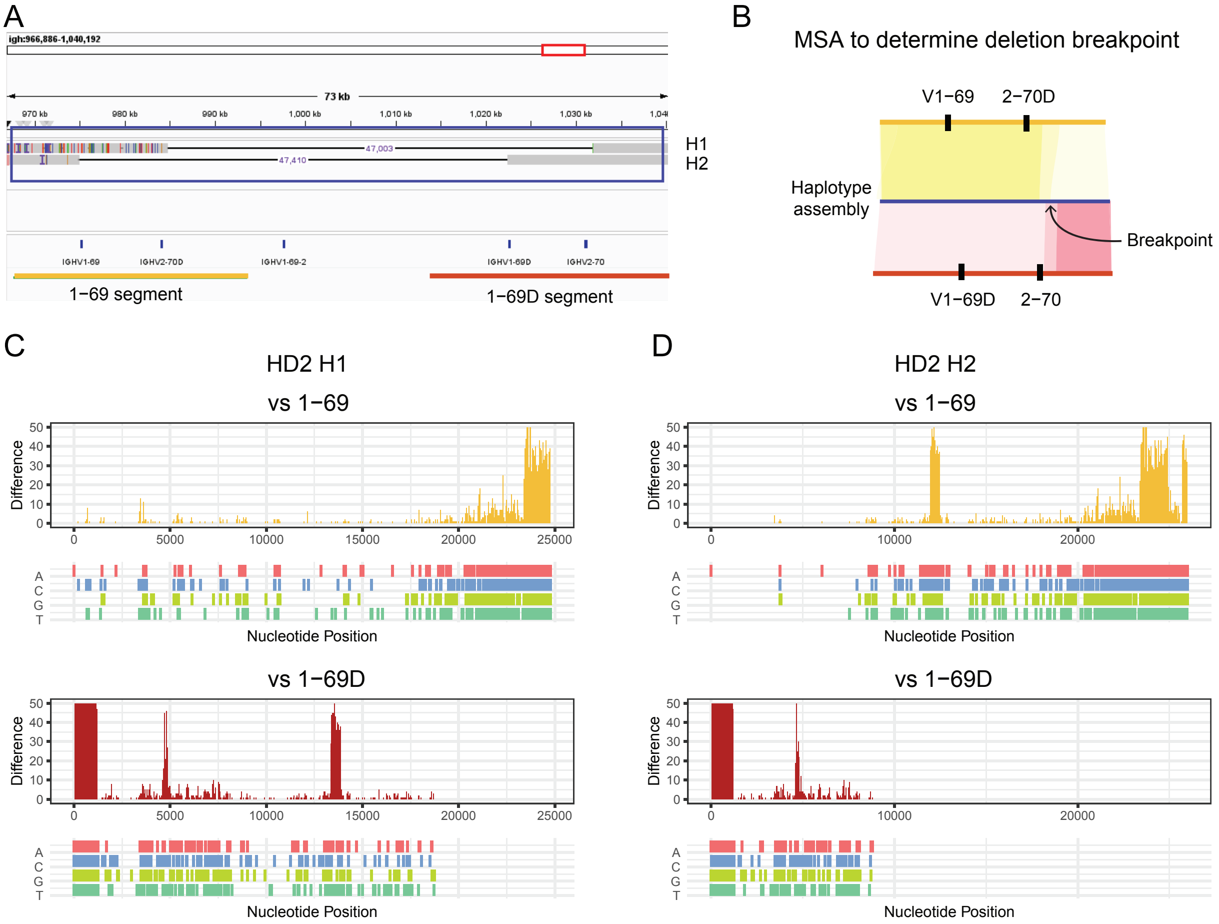


**Figure S2. Breakpoint analysis of deletion haplotypes of IGHV1-69/2-70 in HD2**. (A) IGV window showing Minimap2 alignments of HD2 haplotype assemblies over the IGHV1-69/2-70 region. Blue box shows the segment of the haplotype 1 (H1) and 2 (H2) extracted for multiple sequence alignment (MSA) and breakpoint analysis. Coordinates of segments from IGH reference used for breakpoint analysis highlighted with yellow and red bars in bottom track. (B) Schematic illustrating rationale of breakpoint analysis, higher opacity represents higher identity. Breakpoint is determined to be where the highest identity of HD2 switches from one V1-69/V2-70 segment to the other. (C and D) Nucleotide differences between each V1-69/V2-70 segment and HD2 H1 (C) and H2 (D), shown both as accumulated differences and split by base.

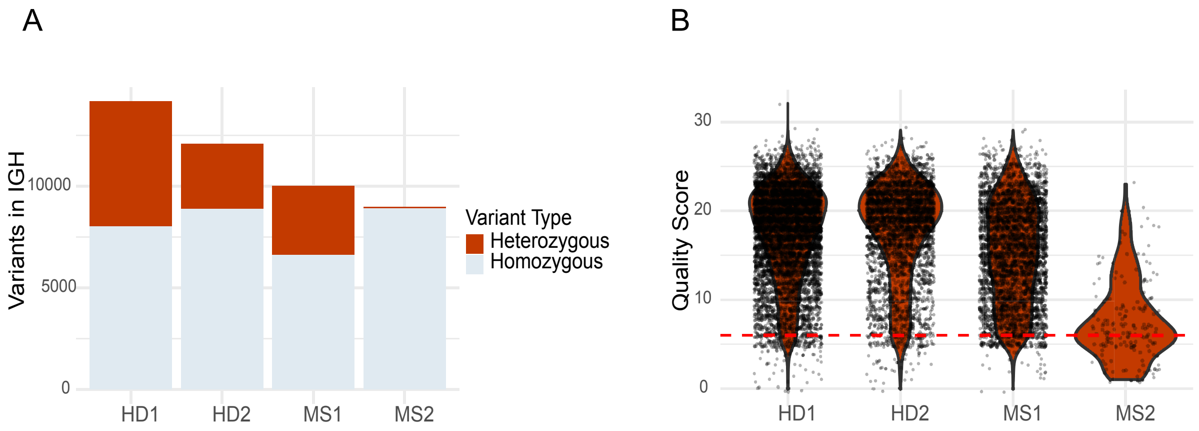


**Figure S3. Analysis of heterozygous variants in IGH, and lack thereof in MS2**. (A) Bar plot showing variants called between the donor's ONT reads and the custom IGH reference by Clair3 during phasing. Variants within the IGH locus that met a Phred quality (Q) score threshold of 6 are stratified by homozygous or heterozygous status. (B) Distribution of variant quality scores for all heterozygous variants in IGH, as determined by Clair3. The stippled line represents the Q=6 threshold required for phasing.

**Table S1. Flagger evaluation of draft and finished haplotype-resolved IGH assemblies for the four donors.** Table shows the number of bases of the dual assembly assigned to each of the four categories.

| | HD1 | | HD2 | | MS1 | | MS2 | |
|---|---|---|---|---|---|---|---|---|
| Donor | Draft | Finished | Draft | Finished | Draft | Finished | Draft | Finished |
| Haploid | 2835084 | 3067029 | 2633025 | 2800672 | 2771599 | 2945333 | 1400097 | 1400097 |
| Collapsed | 51043 | 0 | 125554 | 35052 | 84321 | 0 | 246 | 246 |
| Duplicated | 0 | 0 | 0 | 0 | 0 | 0 | 0 | 0 |
| Error | 0 | 0 | 23793 | 0 | 40884 | 0 | 0 | 0 |

**Table S2. Quantification of IGH genes and alleles.** Summary of genes and alleles annotated in all haplotype assemblies from HD1, HD2, MS1 and MS2.

| Region | Number of genes found | Number of discrete alleles found | Number of alleles with no IMGT MATCH | Number of discrete novel alleles | Number of discrete novel alleles found in VDJbase | Number of discrete alleles not in VDJbase |
|---|---|---|---|---|---|---|
| IGHV | 362 | 138 | 29 | 24 | 21 | 3 |
| IGHD | 177 | 34 | 1 | 1 | 0 | 1 |
| IGHJ | 42 | 7 | 1 | 1 | 0 | 1 |
| IGHC | 69 | 40 | 35 | 25 | 0 | 25 |
| Total | 650 | 219 | 66 | 51 | 21 | 30 |
